# Generalizable links between symptoms of borderline personality disorder and functional connectivity

**DOI:** 10.1101/2023.08.03.551534

**Authors:** Golia Shafiei, Arielle S. Keller, Maxwell Bertolero, Sheila Shanmugan, Dani S. Bassett, Andrew A. Chen, Sydney Covitz, Audrey Houghton, Audrey Luo, Kahini Mehta, Taylor Salo, Russell T. Shinohara, Damien Fair, Michael N. Hallquist, Theodore D. Satterthwaite

## Abstract

**Background:** Symptoms of borderline personality disorder (BPD) often manifest in adolescence, yet the underlying relationship between these debilitating symptoms and the development of functional brain networks is not well understood. Here we aimed to investigate how multivariate patterns of functional connectivity are associated with symptoms of BPD in a large sample of young adults and adolescents.

**Methods:** We used high-quality functional Magnetic Resonance Imaging (fMRI) data from young adults from the Human Connectome Project: Young Adults (HCP-YA; *N* = 870, ages 22-37 years, 457 female) and youth from the Human Connectome Project: Development (HCP-D; *N* = 223, age range 16-21 years, 121 female). A previously validated BPD proxy score was derived from the NEO Five Factor Inventory (NEO-FFI). A ridge regression model with 10-fold cross-validation and nested hyperparameter tuning was trained and tested in HCP-YA to predict BPD scores in unseen data from regional functional connectivity, while controlling for in-scanner motion, age, and sex. The trained model was further tested on data from HCP-D without further tuning. Finally, we tested how the connectivity patterns associated with BPD aligned with agerelated changes in connectivity.

**Results:** Multivariate functional connectivity patterns significantly predicted out-of-sample BPD proxy scores in unseen data in both young adults (HCP-YA; *p*_perm_ = 0.001) and older adolescents (HCP-D; *p*_perm_ = 0.001). Predictive capacity of regions was heterogeneous; the most predictive regions were found in functional systems relevant for emotion regulation and executive function, including the ventral attention network. Finally, regional functional connectivity patterns that predicted BPD proxy scores aligned with those associated with development in youth.

**Conclusion:** Individual differences in functional connectivity in developmentally-sensitive regions are associated with the symptoms of BPD.

## Introduction

Borderline personality disorder (BPD) is a major mental illness that affects 0.7% to 2.7% of adults in the US (1). Individuals diagnosed with BPD experience sudden shifts in mood and struggle to maintain stable interpersonal relationships. BPD is further characterized by impulsivity, suicidality, self-harm, feelings of emptiness, intense anxiety and stress, and dissociative symptoms (1, 2). BPD is also associated with high rates of death by suicide (4%) compared to other mental illness (1, 3-5). BPD and other personality disorders are typically diagnosed in adulthood, but recognizable symptoms often manifest in adolescence (6). Despite their significance, symptoms of BPD are typically not studied in youth samples and the relevant underlying developmental neurobiology remains sparsely explored. Addressing this gap in knowledge is of particular importance given that other major mental illnesses that emerge in adolescence or young adulthood are increasingly understood as disorders of brain development (7).

Previous studies have investigated the link between symptoms of BPD and both brain function and structure using magnetic resonance imaging (MRI) (3, 5, 8, 9), but have yielded inconsistent findings. Studies using resting-state fMRI have reported altered functional connectivity in BPD patients relative to healthy controls in networks associated with emotional processing and executive control (8, 10, 11). Altered functional connectivity has been reported in frontomedial, frontotemporal and limbic regions (12, 13), the fronto-parietal network (10, 14), the default mode network (e.g. posterior cingulate and precuneus (10, 13, 15)), and the salience network (e.g. insula and anterior cingulate cortex (10, 12-14)). More generally, a theoretical perspective on the involvement of frontolimbic circuits in BPD suggests that deficits in the inhibitory function of these regions on circuits associated with social cognition and selfregulation results in emotional dysregulation and behavioral dyscontrol in BPD (16, 17). Although functional alterations in these regions may partly explain the disruptions in emotion and regulatory control processes (e.g., impulsivity) common in BPD, some studies have reported no significant differences in neuroimaging data in patients versus healthy controls (18, 19). These inconsistencies may in part be due to the heterogeneity in BPD populations. However, interacting methodological factors—in particular, small samples—may also be the source of such discrepancies (20).

Prior neuroimaging studies of BPD have mainly used case-control designs with small samples of patients with diagnosed BPD. While such designs can be incredibly valuable and are ultimately essential for clinical translation, the small size of most case-control designs inevitably reduces the replicability and generalizability of the results. Indeed, there is growing evidence that large samples, multivariate models, and out-of-sample testing on unseen data are critical to identify replicable and generalizable brain-behavior associations (20-22). As suggested in part by the Research Domain Criteria (RDoC) (7) and the Hierarchical Taxonomy of Psychopathology (HiTOP) (23) frameworks, one alternative to small case-control designs in psychiatry are dimensional studies of a clinically-relevant construct in larger samples. Furthermore, this perspective accords with overwhelming evidence that BPD symptoms vary dimensionally, with functional impairment scaling with symptom severity (23-27). Although dimensional self-report measures of BPD have been developed (28), substantial evidence also supports the mapping between Big 5 personality trait measures and personality disorder constructs (29). Indeed, Few and colleagues (30) recently developed and validated a trait-based measure of BPD derived from self-reported Big 5 personality traits on the NEO-Five Factor Inventory, which has been collected widely in larger population surveys and clinical samples. The use of such a proxy measure allows the field to leverage existing large-scale data resources with high-quality neuroimaging data to study BPD (19).

Here we aimed to investigate how multivariate patterns of functional connectivity relate to symptoms of BPD in young adults and adolescents using large-scale publicly available datasets. Specifically, we used functional MRI (fMRI) data from two large public datasets to characterize functional connectivity in large samples of adolescents and young adults. We then used machine learning with rigorous cross-validation to predict symptoms of BPD in unseen data from regional patterns of functional connectivity. Finally, to contextualize these results in a developmental framework, we evaluated whether the connectivity patterns that best predicted BPD aligned with age-related changes of functional connectivity in youth.

## Methods and Materials

We used functional connectivity networks from two large-scale, publicly available datasets—the Human Connectome Project - Young Adult (HCP-YA (31)) and the Human Connectome Project - Development (HCP-D (32))—to predict individual differences in BPD symptoms as measured by a validated trait-based BPD proxy score. We note that here the term “predict” refers to contemporaneous association between BPD and functional connectivity in *unseen* data rather than prospective prediction of BPD. As described below, to investigate the link between regional functional connectivity and BPD scores in the adult data from HCP-YA, a multivariate linear ridge regression model was first trained for each brain region, predicting participants’ BPD scores (i.e., dependent variable) from multivariate functional connectivity patterns (i.e., independent predictor). The trained model was then tested on unseen data using cross-validation. Next, as a strong test of the generalizability of our model, we applied a fully trained model from HCP-YA to data from HCP-D without further training or tuning, testing the model on unseen data (i.e., HCP-D was not part of the training). Finally, we compared the degree to which connectivity patterns that best predicted BPD in this model corresponded to functional connectivity patterns that displayed the greatest developmental effects in HCP-D. Throughout, we performed multiple sensitivity analyses to assess the robustness of our findings.

### Participants

#### Young adult sample

Resting-state and task functional Magnetic Resonance Imaging (fMRI) data from healthy young adults were obtained from the Human Connectome Project–Young Adults (HCP-YA (31); *N* = 870, ages 22-37 years, 457 female and 413 male). All imaging data were collected on a customized 3 Tesla scanner at Washington University (WashU). Data included 1 hour of resting-state fMRI scans acquired over 2 days (2 scans along opposing R/L and L/R phase encoding directions per day; each about 15 minutes long) and 1 hour of task-fMRI data over 2 days (one scan of 30 minutes long per day, where each task was run twice with opposing R/L and L/R phase encoding directions). All functional images were acquired with high spatial and temporal resolution (8x multiband acquisition, 2 mm^3^, TR=720 ms, TE=33.1 ms, flip angle=52°).

#### Developmental sample

The developmental sample included resting-state and task fMRI data from the Human Connectome Project–Development 2.0 Release (HCP-D (32); *N* = 610, age range 5.6-21.9 years, 331 female and 279 male). Imaging data were obtained on 3 Tesla Siemens Prisma platforms at four sites. Participants who were at least 8 years old completed 26 minutes of resting-state fMRI scans in four runs, whereas the scan time for younger participants (5-7 years old) were reduced to 21 minutes in total across all runs. The data also included 8 minutes of task-fMRI data across two runs. All functional images were acquired with high spatial and temporal resolution (8x multiband acquisition, 2 mm^3^, TR=800 ms, TE=37 ms, flip angle=52°).

### Image processing

The two datasets (HCP-YA and HCP-D) were preprocessed with similar image processing pipelines as described below.

#### Young adult sample

All images were processed with HCP minimal preprocessing pipelines (33). The minimally preprocessed HCP outputs were then post-processed using the eXtensible Connectivity Pipelines-DCAN collaborative (XCP-D (34)). The preprocessing included gradient distortion correction, field-map distortion correction, boundary-based registration, bias field correction, and whole brain intensity normalization. Functional timeseries were then projected to the fsLR cortical surface and smoothed with a 2mm FWHM Gaussian kernel. Post-processing included despiking (3dDespike as implemented by AFNI (35, 36)) and confound regression of six motion parameters (3 translation (x,y,z) and 3 rotational (roll, pitch, yaw) motion measurements), the mean surface timeseries signal as the global signal, the mean white matter, the mean cerebrospinal fluid signal with their temporal derivatives, and the quadratic expansion of the six motion parameters, tissues signals, and their temporal derivatives. Residual timeseries from this regression were then band-pass filtered between 0.01 and 0.08 Hz. A summary measure of in-scanner motion was estimated as framewise displacement, and high-quality data with low mean framewise displacement (mean [FD] <0.2mm) were retained for subsequent analyses. Finally, the preprocessed time-series were parcellated into 400 cortical regions using the Schaefer-400 atlas (37). The parcellated time-series were used to construct functional connectivity matrices as Pearson correlation coefficients between pairs of regional time-series for each participant. We also used a lower resolution parcellation with 200 cortical regions (i.e., the Schaefer-200 atlas) for sensitivity analysis.

#### Developmental sample

The developmental data (HCP-D) was preprocessed with the ABCD-BIDS pipeline (38), which is an updated version of HCP minimal preprocessing pipeline with a few differences and additional steps. Specifically, ABCD-BIDS uses the Advanced Normalization Tools (ANTs) for denoising, bias field correction, and nonlinear registration, which improves the registration performance compared to other methods (4, 39). Moreover, the motion estimation process in ABCD-BIDS provides an estimate of framewise displacement after accounting for respiratory effects on magnetic field changes (40). In addition, given that the HCP-D data was a higher motion sample compared to HCP-YA, scrubbing was used to remove high-motion time frames that had larger than 0.2 mm framewise displacement estimates (41). The rest of the image processing steps were the same as described above for HCP-YA.

### Assessment of BPD-spectrum symptoms

Targeted measures of BPD symptoms are usually administered only in focal studies of BPD in adults and are not available in large-scale neuroimaging studies of youth. To obtain a proxy measure of BPD-spectrum symptoms in HCP-YA and HCP-D, we used a previously-validated proxy measure of BPD (30) that has been previously used to investigate BPD-spectrum symptoms in large imaging datasets such as the HCP-YA (19). This proxy measure estimates BPD symptom scores using 24 items from a widely-used personality assessment instrument, NEO Five Factor Inventory (NEO-FFI). Few and colleagues developed and validated this trait-based BPD proxy score across multiple datasets, comparing the BPD proxy score with explicit measures of BPD in both clinical BPD samples and the broader population (30). Given that the NEO-FFI is available in most large-scale datasets, it provides a useful and scalable route to study BPD-relevant symptoms in large-scale data. We used the NEO-FFI instrument to estimate a BPD proxy score for each participant for both young adults (HCP-YA) and adolescents (HCP-D) as previously described (30) (**Table S1**). Note that NEO-FFI was available for all participants in HCP-YA data, whereas it was only available for participants over age 16 in HCP-D data (*N* = 223, 121 female and 102 male).

### Multivariate analyses

Prior work has shown that identifying reliable and generalizable brain-behavior associations requires out-of-sample testing, and that reliability is enhanced by multivariate analysis approaches (20, 21). Accordingly, here we used a machine learning approach to predict BPD proxy scores from multivariate regional functional connectivity patterns. Specifically, we used linear ridge regression modeling as implemented in Scikit-Learn (42). We trained a separate linear ridge regression model for each brain region, predicting each participant’s BPD proxy score from the region’s functional connectivity profile (i.e., all connections between a given region and all other regions). Hence, the dependent variable was the participant’s BPD score and the independent predictor was a row of their functional connectivity matrix (**Figure 1**). This analysis yielded an estimate of the association between BPD symptoms and the functional connectivity profile of each brain region. As described below, all models included covariates for age, sex, and in-scanner motion (mean FD).

**Figure 1.**
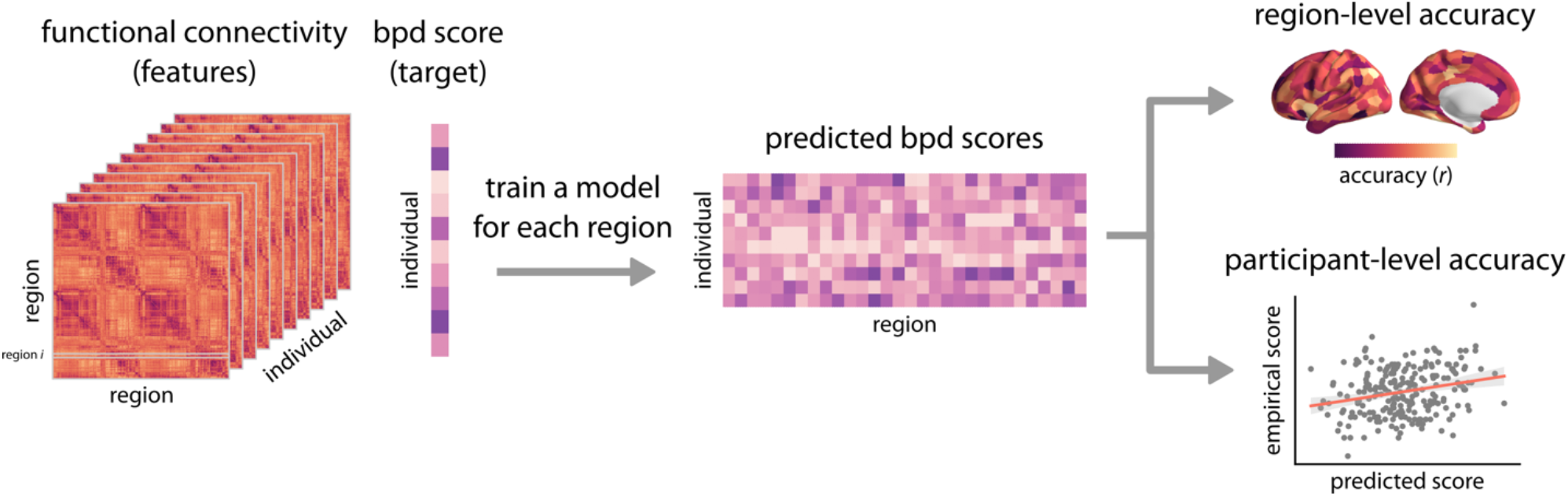
Predicting BPD proxy scores from multivariate functional connectivity patterns. Functional connectivity data (i.e. features) were used to predict BPD proxy scores (i.e., target) in young adults and adolescents. A separate linear ridge regression model was trained using a given region’s functional connectivity profile (e.g., the *i*-th row of the connectivity matrix corresponding to the connections between region *i* and all other regions). The regional trained models were applied to the held-out test data to obtain predicted BPD scores for each individual and each region. Region-level accuracy was estimated as the Pearson correlation coefficient *r* between the empirical and predicted BPD scores for each model. Participant-level accuracy was estimated as the Pearson correlation coefficient *r* between the empirical and average predicted BPD scores across regional models.

We first trained and tested the ridge regression models on the HCP-YA data using 10-fold cross-validation. For each split, we performed nested cross-validation and hyperparameter tuning (regularization parameter = [0.1, 1.0, 10]) using the training set. To avoid leakage across training and test sets when correcting for covariates, we first used a linear regression model to remove covariate effects from connectivity data in the training set. We then used the same linear regression model (without any additional adjustments) to correct for covariates in the test set. Note that the linear regression model coefficients were defined only based on the training set, avoiding any information leakage in the test data. This procedure was repeated separately for each train and test split. Following the covariate correction process, the ridge regression model trained on the training set was applied to the held-out test set to predict the BPD proxy scores in the *unseen data*. To assess the final out-of-sample model performance of a given brain region in HCP-YA, we pooled the test set predicted scores across all 10 folds and estimated the model performance as the Pearson correlation coefficient *r* between the empirical and predicted BPD proxy scores. To obtain the participant-level model performance, we calculated the average predicted score across brain regions for each participant. The model performance was then estimated as the correlation coefficient between the empirical and average predicted scores across regional models. We evaluated the significance of predictive performance using permutation testing: we generated a null distribution of predicted scores for each split using 1,000 permutations by randomly shuffling the dependent variable (i.e., BPD scores) and repeating the analysis. The model performance was assessed as described above. This procedure yielded null distributions of participant-level and region-level model performance.

To assess system-specific model performance, we calculated the average model performance in the seven resting-state functional networks (i.e., intrinsic networks) as defined by Yeo and colleagues (43). We then performed a network enrichment analysis using spatial autocorrelation-preserving null models (i.e., “spin” tests) (44, 45). Specifically, we generated null brain maps of regional model performance with disrupted topographic patterns but preserved spatial autocorrelation by applying 10,000 randomly-sampled rotations to the spherical projections of the data (44). The rotations were applied to one hemisphere and then mirrored to the other hemisphere. We calculated the average model performance for each intrinsic network in the null brain maps, generating a null distribution of average model performance per intrinsic network.

Given that multivariate analysis along with large samples and out-of-sample testing are essential for generalizable studies of brain-behavior associations, we next aimed to directly evaluate the generalizability of our approach. We selected HCP-D data since it is a large dataset with a younger age range and is acquired using different sequences and at different scanning sites, making it a good candidate to test the generalizability of our approach and predictive models. We used all the data from HCP-YA to train a ridge regression model, and then applied the trained regional models to the completely unseen HCP-D data without any additional tuning. Model performance and significance were evaluated as described above.

### Functional connectivity age effects

To assess whether the association between functional connectivity and BPD proxy scores reflects any age-related effects on functional connectivity, we estimated how regional functional connectivity profiles evolve during development. We used a linear ridge regression model for each region to predict participant age at the time of the interview from the region’s functional connectivity profile in the full developmental sample (HCP-D; *N* = 610), while controlling for in-scanner motion and sex as above. As in prior analyses, the models were trained and tested using a 10-fold cross-validation analysis. The regional age prediction accuracies were then compared with regional BPD proxy score prediction accuracies for the HCP-D dataset. We performed 10,000 spin permutations to compare the topographic patterns across the cortex using spatial autocorrelation-preserving null models generated as described above (44, 45). This procedure allowed us to evaluate if the regions whose functional connectivity profiles best predicted BPD symptoms co-localized with regions whose connectivity varied the most with brain development.

## Code and data availability

Data used in the present study were obtained from publicly available HCP-YA and HCP-D datasets (31, 32). Code used to conduct the analyses reported in this study is available on GitHub (https://github.com/PennLINC/borderline).

## Results

### BPD score distribution

BPD proxy score was estimated from 24 NEO-FFI items in both young adults (HCP-YA: *α* = 0.79, mean = 0.90, SD = 0.39) and adolescents (HCP-D: *α* = 0.80, mean = 1.17, SD = 0.43). Summary measures from BPD score distributions and sample descriptions are included in **Table 1** for both datasets. The BPD score distribution in both samples was comparable to the BPD scores reported by Few and colleagues (30) in college students (Sample 1: *α* = 0.78, mean = 1.76, SD = 0.44; Sample 2: *α* = 0.73, mean = 1.83, SD = 0.43) and a general population of young adults (*α* = 0.81, mean = 1.52, SD = 0.43). Note that Few and colleagues reported higher BPD scores (*α* = 0.67, mean = 2.29, SD = 0.39) for psychiatric outpatient adult populations (including patients with BPD).

**Table 1.** BPD score distribution. Sample descriptions and BPD score distributions (range, mean, and standard deviation (SD)) are provided. Note that the BPD proxy score was only estimated for a subset of individuals in HCP-D since the NEO-FFI questionnaire was only available for individuals over age 16. Internal consistency was estimated using Cronbach’s *α*.

|  | HCP-YA<br>(*N* = 870) | HCP-D (with NEO-FFI)<br>(*N* = 223) | HCP-D (full sample)<br>(*N* = 610) |
| --- | --- | --- | --- |
| Age<br>(range, mean, SD) | [22.0, 27.0], 28.6, 3.7 | [16.0, 21.9], 19.0, 1.8 | [5.6, 21.9], 14.6, 4.0 |
| Sex<br>(female, male) | 457, 413 | 121, 102 | 331, 279 |
| BPD score<br>(range, mean, SD) | [-0.17, 2.30], 0.90, 0.39 | [0, 2.75], 1.17, 0.43 | n/a |
| Reliability coefficient<br>(Cronbach's $\alpha$, CI) | 0.79, [0.77, 0.81] | 0.80, [0.78, 0.82] | n/a |

### Functional connectivity predicts symptoms of BPD

We used a multivariate linear ridge regression model to predict BPD proxy scores from a given brain region’s functional connectivity profile (**Figure 1**). The regional models were initially trained on the data from healthy young adults (HCP-YA) and tested on unseen data from the same cohort using cross-validation. We found that multivariate patterns of functional connectivity significantly predicted dimensional BPD symptoms (**Figure 2a;** *r* = 0.14, *p*_perm_ = 0.001). These findings demonstrate that multivariate functional connectivity patterns are indeed linked to symptoms of BPD in young adults. We further found that there was substantial heterogeneity in how well the functional connectivity profile of each region predicted BPD symptoms (**Figure 2b**; see regional maps thresholded based on permutation tests in **Figure S1**). To investigate whether the regional prediction accuracy was more pronounced in specific functional systems, we estimated the average prediction accuracy for the 7 intrinsic functional networks defined by Yeo and colleagues (43) (**Figure 2c**). We found that the prediction accuracy was highest for the fronto-parietal (*p*_spin_ = 0.021; FDR-corrected) and ventral attention (*p*_spin_ = 0.007; FDR-corrected) networks, suggesting a link between BPD symptoms and systems involved in emotion regulation and executive function.

**Figure 2.**
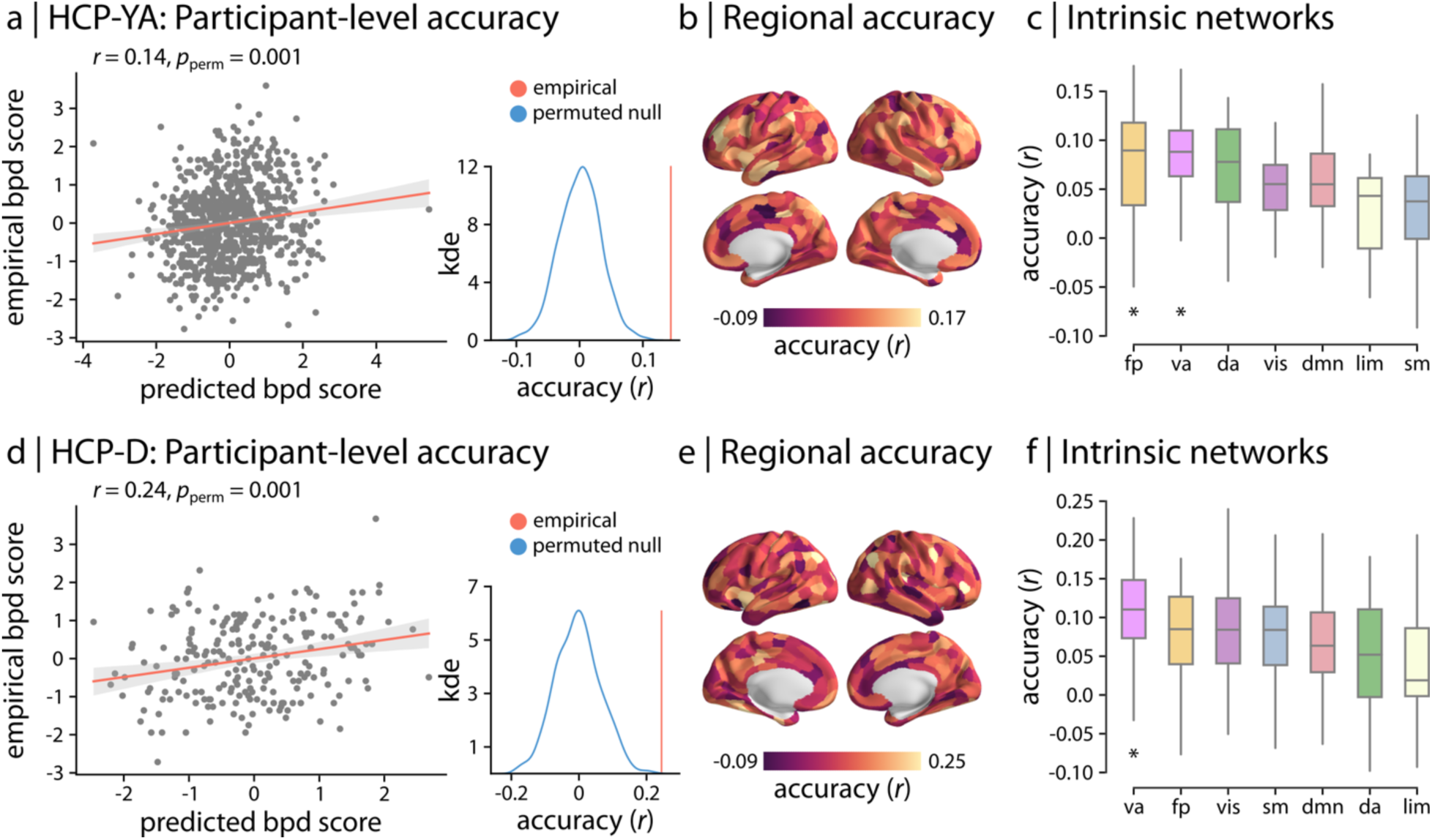
Functional connectivity predicts BPD scores in young adults and adolescents. Regional linear ridge regression models were used to predict BPD proxy scores from multivariate functional connectivity patterns in (a-c) healthy young adults from the Human Connectome Project (HCP-YA) data and (d-f) adolescents from the Human Connectome Project - Development (HCP-D). The model performance was assessed as the Pearson correlation coefficient *r* between the empirical and predicted scores. Participant-level model performance is depicted as scatter plots, demonstrating the relationship between the empirical and average BPD scores across regional models (HCP-YA: *r* = 0.14, 95% CI = [0.08 0.21]; HCP-D: *r* = 0.24, 95% CI = [0.12 0.37]). Each point in the scatter plot represents a participant. The participant-level accuracy *r* (red vertical line) is then compared with a null distribution of accuracies (blue distribution) obtained from 1,000 permutation tests, randomly shuffling the samples. Regional out-of-sample model performance is depicted across the cortex for both cohorts (Schaefer-400 atlas; 99% confidence intervals) (see regional maps thresholded based on permutation tests in **Figure S1**). Finally, the average functional system-level prediction accuracy was estimated for the 7 intrinsic functional networks defined by Yeo and colleagues (32). Asterisk denotes significant system-level prediction accuracy based on 10,000 spatial autocorrelation-preserving null models (*p*_spin_ < 0.05; FDR-corrected). Significant system-level accuracy was observed in fronto-parietal (*p*_spin_ = 0.021; FDR-corrected) and ventral attention (*p*_spin_ = 0.007; FDR-corrected) networks for HCP-YA and ventral attention network (*p*_spin_ = 0.0001; FDR-corrected) for HCP-D. Intrinsic networks: vis = visual; sm = somatomotor; da = dorsal attention; va = ventral attention; lim = limbic; fp = fronto-parietal; dmn = default mode.

### BPD symptoms are linked to functional connectivity in adolescents

To examine whether the link between functional connectivity and BPD symptoms identified in early adulthood generalizes to late adolescence, we used the previously trained regional models to predict BPD proxy scores in HCP-D. Note that the regional models were trained on HCP-YA data *only* and the trained models were applied to HCP-D; as such, the HCP-D data were completely unseen by the models during training. Consistent with the reported findings in young adults, we found that regional functional connectivity profiles significantly predict BPD proxy scores in adolescents (**Figure 2d;** *r* = 0.24, *p*_perm_ = 0.001). The fact that the model constructed in young adults significantly predicted BPD symptoms in an unseen sample of older adolescents suggests that multivariate connectivity patterns are generalizably linked to BPD symptoms across ages and multiple datasets. Notably, regional patterns showed similar heterogeneity as in the adult data (**Figure 2e;** see correspondence between regional accuracies across samples in **Figure S2**), with the highest prediction accuracy observed in ventral attention network (FDR-corrected *p*_spin_ = 0.0001; **Figure 2f**). Together, these results suggest that individual differences in brain systems important for emotion regulation and executive function are linked to BPD symptoms in both young adults and adolescents.

### Regions that predict BPD symptoms also display developmental effects

Next, we sought to situate our findings in the context of brain development. Specifically, we evaluated whether the regions that were most strongly linked to BPD symptoms also were those that displayed age-related changes in connectivity during development in youth. To quantify the developmental effects in functional networks, we used regional patterns of functional connectivity to predict the age of unseen HCP-D participants. We found that functional connectivity was associated with age, but that prediction accuracy varied significantly across the cortex (**Figure 3a**). To directly examine if these age-related changes in connectivity profiles aligned with regions associated with BPD symptoms, we compared the cortical distributions of age prediction accuracy and BPD proxy score prediction accuracy in the developmental sample (**Figure 3b**). Permutation testing with spatial autocorrelation-preserving null models (i.e., “spin” tests) revealed a significant association between cortical distribution of age and BPD proxy score prediction accuracy (**Figure 3b**; *r* = 0.20, *p*_spin_ = 0.001). This finding suggests that regions associated with BPD symptoms are also those that undergo greater age-related changes in functional connectivity in youth.

**Figure 3.**
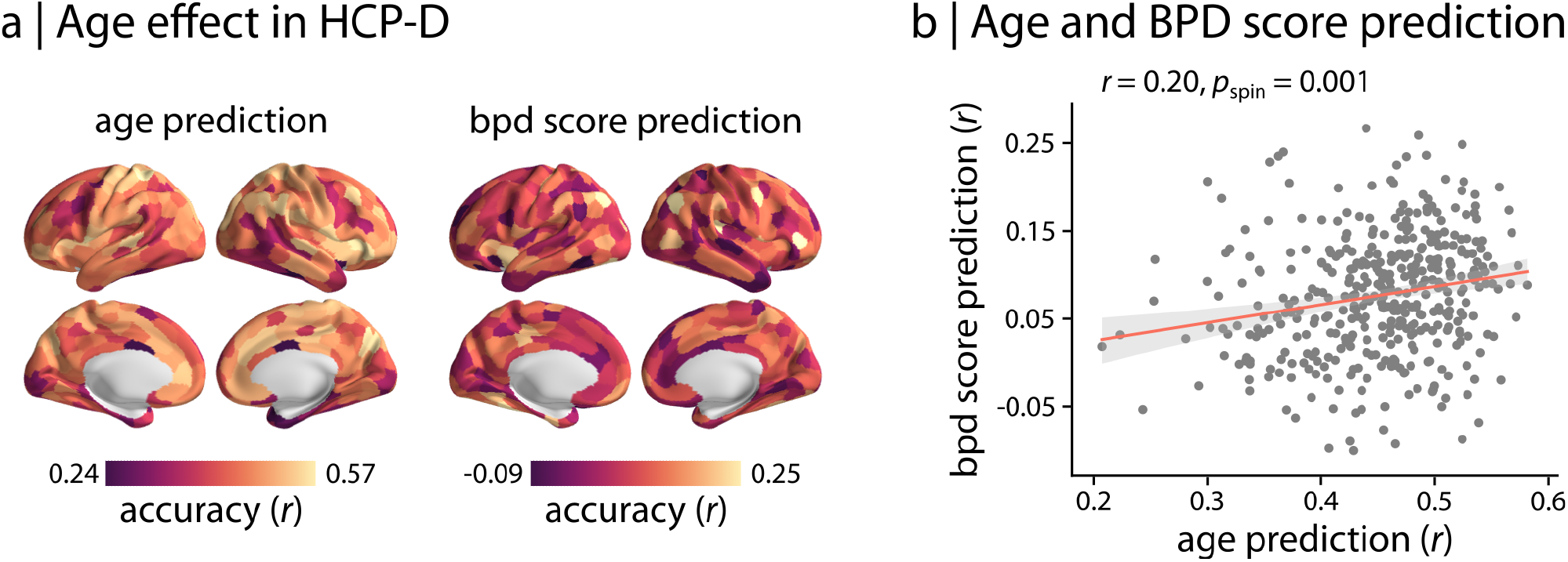
Regional predictive capacity aligns with developmental changes in functional connectivity. (a) Regional functional connectivity profiles were used to predict participants’ age in the developmental sample (HCP-D). The age prediction accuracy *r* is depicted across the cortex along with the previously obtained BPD proxy score prediction accuracy (Schaefer-400 atlas; 99% confidence intervals). (b) Topographic patterns of BPD score prediction and age prediction were compared using the Pearson correlation coefficient *r* and 10,000 spatial autocorrelation-preserving null models (i.e., “spin” tests). Each point in the scatter plot corresponds to a brain region.

### Sensitivity analysis provides convergent results

As a final step, to evaluate if our findings were affected by specific analytical choices, we performed multiple sensitivity analyses. First, we repeated the analyses using only resting-state fMRI data, rather than the concatenated task- and rest-fMRI scans used in the original analysis. Specifically, we used functional connectivity networks from only resting-state fMRI data to predict BPD proxy scores in both young adults (HCP-YA; **Figure S3a**) and adolescents (HCP-D; **Figure S3b**). The results were consistent in both samples. Next, to ensure that the reported findings were robust to the parcellation resolution chosen, we repeated all the analyses with a different parcellation resolution (200 rather than 400 regions). Again, results were consistent for both HCP-YA (**Figure S4a**) and HCP-D (**Figure S4b**). Finally, to verify that the findings were not influenced by scanning-site differences in HCP-D, we used CovBat-GAM (46-49) to harmonize functional connectivity data across sites and repeated the analyses with harmonized data. The results were consistent with the original findings (**Figure S5**). These sensitivity analyses— combined with the generalizability of results across samples—bolsters confidence in the reported findings.

## Discussion

To our knowledge, this is the largest functional neuroimaging study of BPD symptoms in adolescence and young adulthood. We found three main results. First, functional connectivity significantly predicted symptoms of BPD in large samples of both young adults (HCP-YA) and adolescents (HCP-D). Second, the predictive capacity was heterogeneous across the cortex, such that the most predictive regions were found in functional systems relevant for emotion regulation and executive function. Finally, we found that regions associated with BPD symptoms co-localized with regions with prominent age-related changes in connectivity in youth.

Previous studies have sought to associate symptoms of BPD with diverse functional and structural neuroimaging markers (3, 8). The findings from these studies have varied considerably and as yet there is no consensus regarding how differences in brain function are linked to BPD symptoms. However, one relatively consistent finding across studies is the presence of altered structural and functional patterns in fronto-limbic networks implicated in emotion regulation and cognitive control (8, 10-14, 50). One likely cause of the heterogeneity in the existing literature is the relatively small samples studied, hampering efforts to identify reliable and replicable neurobiological signatures of BPD symptoms (3).

It should be noted that such challenges are hardly unique to BPD research; identifying reliable, replicable, and generalizable neuroimaging indices of psychopathology remains a major ongoing challenge (20, 22). Recent evidence suggests that large samples, out-of-sample testing, and use of multivariate methods are essential for studies that seek to link brain and behavior (20-22). Out-of-sample testing using rigorous cross-validation analysis along with analyses that assess the generalization of results to unseen, new datasets collected under different conditions are required to properly assess brain-behavior associations. Fortunately, collaborative efforts in collecting and sharing human neuroimaging data with large sample sizes have provided an unprecedented opportunity to examine the brain-behavior association in a systematic and comprehensive manner. Although there has been a growing number of largescale publicly available datasets in healthy populations and clinical cohorts (e.g., ADNI for Alzheimer’s Disease (51); PPMI for Parkinson’s Disease (52); ENIGMA consortium for various neurological and psychiatric diseases and disorders (53)), most neuroimaging studies that are focused on neuropsychiatric brain disorders and diseases have small sample sizes. An alternative approach suggested in part by the RDoC initiative and the HiTOP system is to investigate dimensional variations of clinically-relevant measures in larger non-clinical samples such as HCP (7, 23).

We used a personality trait-based measure to estimate BPD proxy scores in large samples of healthy young adults and adolescents. This allowed us to take advantage of publicly available datasets to study the link between brain function and BPD symptoms. One recent study used a similar approach to investigate the link between structural neuroimaging markers from T1weighted MRI data and BPD symptoms in two large samples (19). However, the study identified no associations between BPD symptoms and the MRI structural markers, including cortical thickness, surface area and subcortical volumes (19). Considered together with our findings, this aligns with the recent evidence indicating that functional brain organization may provide a more sensitive tool than anatomical markers for capturing brain-behavior relationships in some settings (20). Here, we investigated the link between multivariate functional connectivity patterns and BPD symptoms in adolescence and young adulthood. Our findings demonstrated that functional connectivity is indeed associated with BPD proxy scores in both adolescents and young adults. The spatial distribution of regional predictive capacity was heterogeneous across the cortex. Consistent with some studies in the previous literature, the link between functional connectivity and BPD symptoms was more prominent in certain functional systems, such as the ventral attention network (which overlaps with insular cortex and the salience network) and the fronto-parietal network (11, 50). These functional networks are associated with emotion regulation (ventral attention network) and executive function and top-down control (frontoparietal network). The high contribution of ventral attention and fronto-parietal networks to the link between brain function and BPD symptoms may be linked to impairments in emotion regulation and impulse control reported in patients with BPD (1).

Notably, we found that this link between functional connectivity and BPD proxy score was generalizable across datasets of both young adults and adolescents collected at different sites. This finding indicates that although BPD is usually not diagnosed before the age of 18 years (2), individual differences in functional networks linked to BPD may be present earlier in development. This possibility is particularly relevant given the finding that brain regions that undergo the most functional maturation during development are the ones that contribute the most to the link between functional connectivity and BPD symptoms. Together, these findings suggest that—like many other major neuropsychiatric conditions (7, 54)—BPD may be understood in part as a disorder of neurodevelopment. Examining BPD from a neurodevelopmental perspective may accelerate efforts to identify markers of risk for BPD earlier in life and develop personalized interventions before negative outcomes of this disabling disorder accrue.

The findings presented in this work should be considered with respect to several methodological caveats. First, the analyses were not performed in patients diagnosed with BPD. The two datasets used in this study include healthy young adults and typically developing adolescents. The main objective of this study was to conduct a large-scale functional imaging study of BPD symptoms leveraging the large sample sizes of publicly available datasets. Although we used a trait-based BPD proxy score that has been previously validated in multiple datasets including healthy and BPD populations, future work applying this approach to individuals diagnosed with BPD (or individuals with greater levels of BPD symptomatology) using targeted BPD measures is required to further confirm our findings. Second, the NEO-FFI questionnaire, and hence the BPD proxy score used here, was only available in older adolescents and young adults (individuals who were at least 16 years or older). Using this trait-based measure of BPD proxy score made it possible to expand our analysis to older adolescents, which have generally not been included in prior studies of BPD. Moving forward, the link between functional brain organization and BPD symptoms in younger adolescents remains an important area for study, given that recognizable symptoms of BPD often manifest much earlier in life, even as early as 12 years old (1, 2). Third, BPD is a heterogeneous disorder and patients with BPD often suffer from comorbid mental health problems and other psychiatric conditions (1, 5). Further work is required to discover the impact of comorbidity with other mental disorders. Finally, the goal of this study was to reliably identify a link between multivariate patterns of functional connectivity and BPD using large sample and rigorous out-of-sample analysis. Future investigation focusing on specific functional connections and regions-of-interest is essential to identify detailed neural substrates of BPD.

In sum, we demonstrated that multivariate functional connectivity patterns can successfully predict BPD symptoms in unseen data from both young adults and adolescents. Findings also suggest regions whose functional connectivity develops the most in youth align with those associated with BPD, providing new evidence for understanding BPD as a neurodevelopmental disorder. Moving forward, linking within-individual neurodevelopmental trajectories of functional connectivity to the emergence of BPD is an important direction for longitudinal studies.

## Acknowledgments

Funding for this study was provided by the AE foundation. Additional support was provided by the National Institutes of Health: R01MH113550 (TDS & DSB), R01MH120482 (TDS), R01EB022573 (TDS), R37MH125829 (TDS & DAF), R01MH123550 (RTS), R01MH112847 (RTS & TDS), R01MH123563 (RTS). GS was supported by a postdoctoral fellowship from the Canadian Institutes of Health Research (CIHR). Additional support was provided by the Penn-CHOP Lifespan Brain Institute.

---

**Figure S1.**
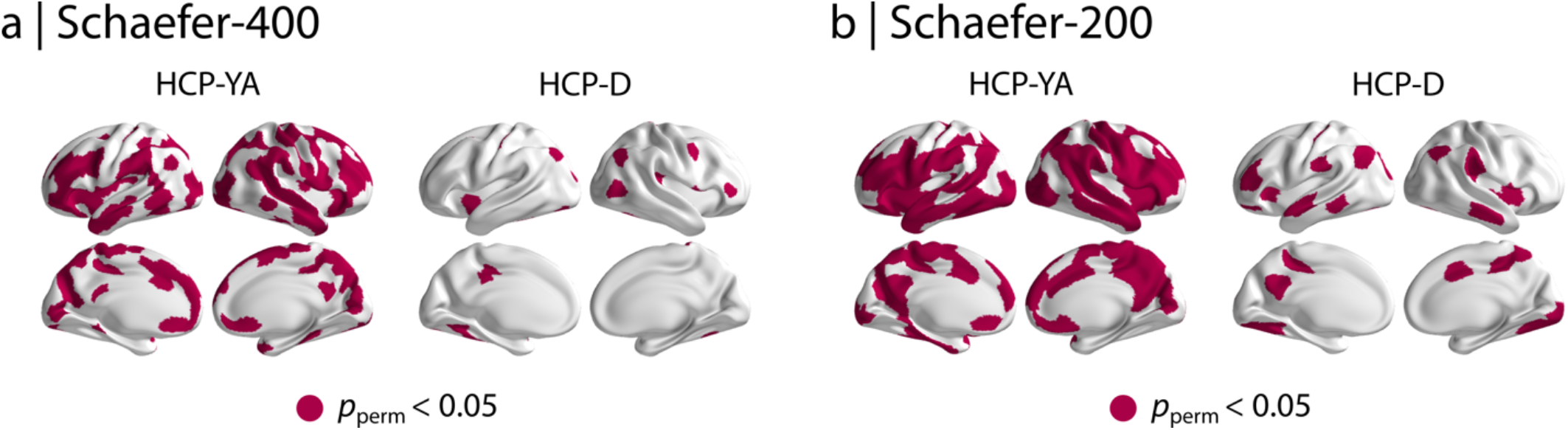
Regional predictive capacity thresholded based on permutation tests. We generated a null distribution of predicted scores for each regional model using 1,000 permutations by randomly shuffling the dependent variable (i.e., BPD scores) and re-calculating the model performance for each permutation. Regional maps were then thresholded based on permutation tests following correction for multiple comparisons (*p*_perm_ < 0.05; FDR-corrected).

**Figure S2.**
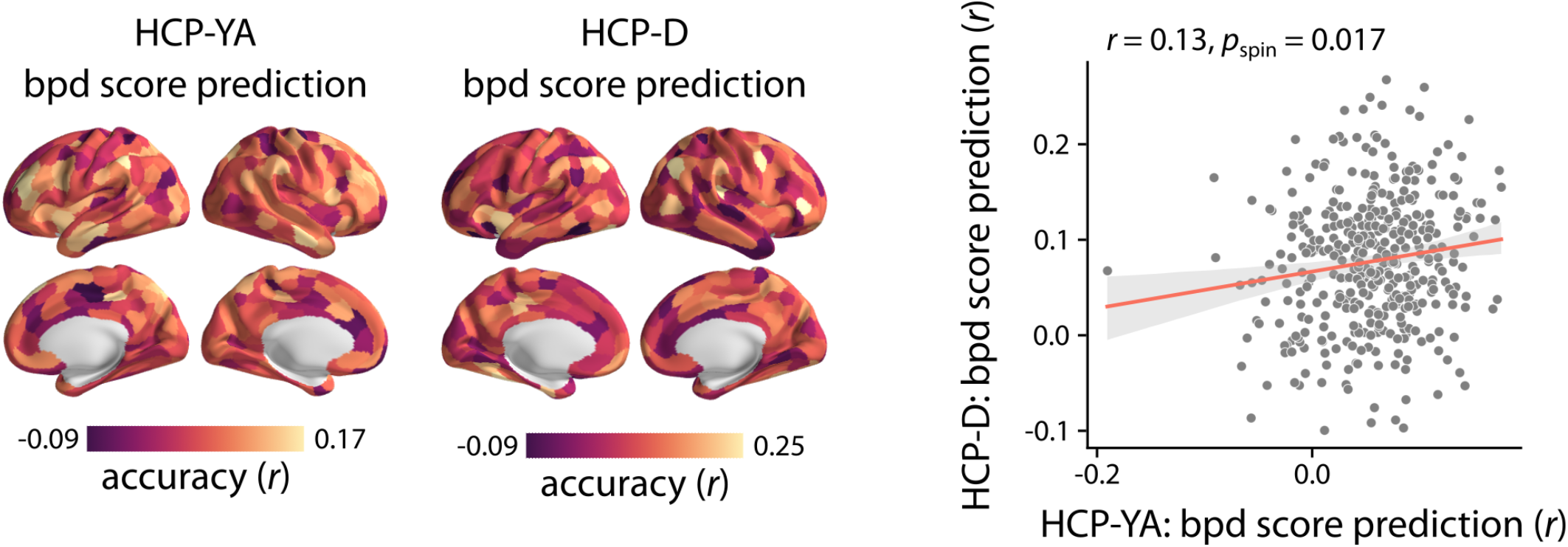
Regional predictive capacity is consistent in adulthood and adolescence. Spatial distributions of BPD score predictions in adulthood (HCP-YA; **Figure 2a**) and adolescence (HCP-D; **Figure 2b**) were compared using Pearson correlation coefficient *r* and 10,000 spatial autocorrelation-preserving null models (i.e., “spin” tests). Each point in the scatter plot corresponds to a brain region.

**Figure S3.**
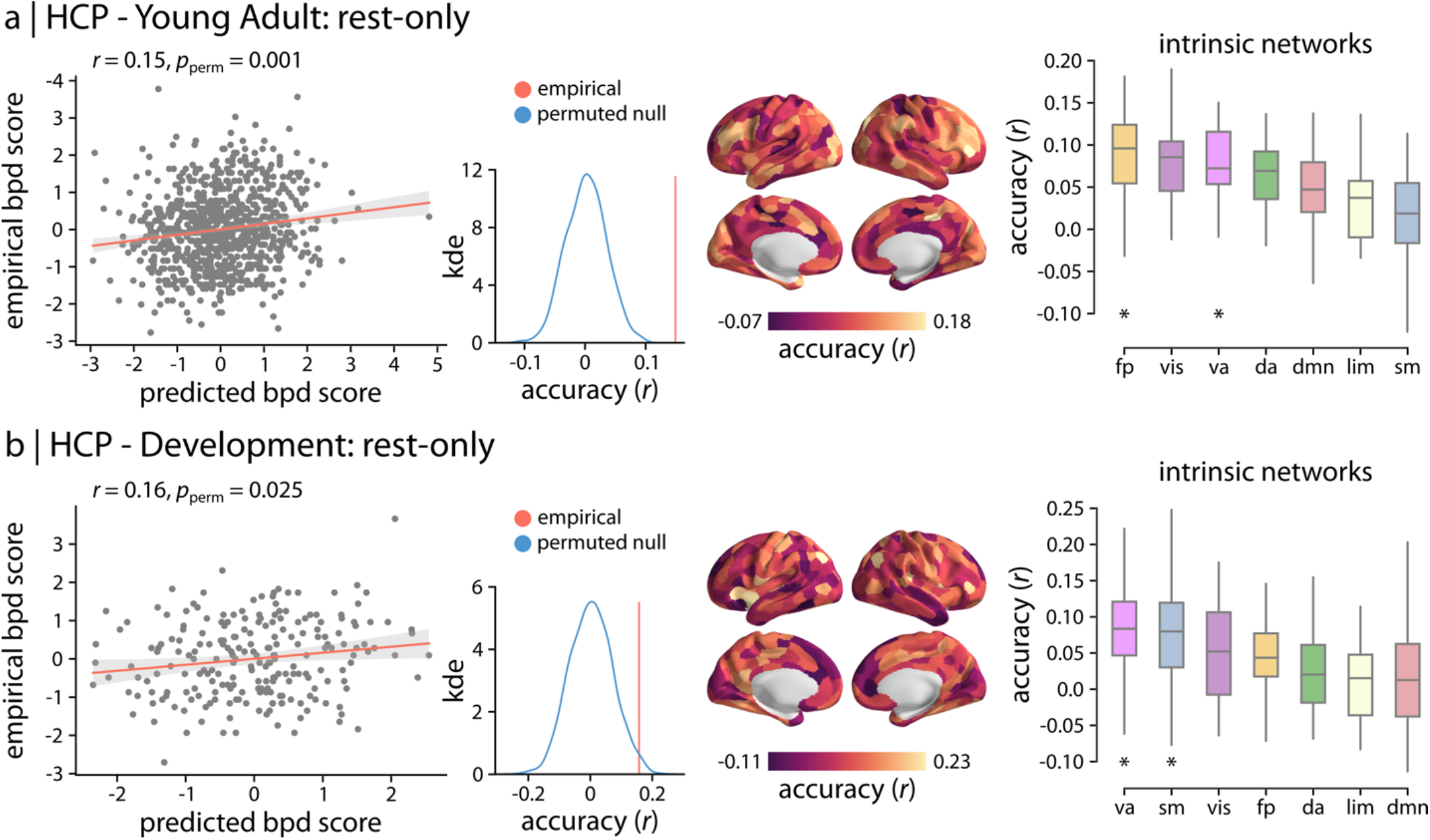
Sensitivity analysis with resting-state fMRI only. Only the resting-state fMRI data were used to predict BPD proxy scores, rather than the concatenated rest- and task-fMRI data used in the original analysis (Figure 2). The results are depicted for (a) healthy young adults from the Human Connectome Project (HCP-YA) data and (b) adolescents from the Human Connectome Project - Development (HCP-D). The participant-level prediction accuracies are shown using the scatter plots (HCP-YA: *r* = 0.15, 95% CI = [0.08 0.21]; HCP-D: *r* = 0.16, 95% CI = [0.02 0.28]). Similar to the original analysis, the results were compared with null distributions of accuracies obtained from permutation tests. Region-level accuracies are also shown across the cortex (Schaefer-400 atlas; 99% confidence intervals). Finally, the average functional system-level prediction accuracy was estimated for the 7 intrinsic functional networks. Asterisk denotes significant system-level prediction accuracy based on 10,000 spatial autocorrelation-preserving null models (*p*_spin_ < 0.05; FDR-corrected). Significant system-level accuracy was observed in fronto-parietal (*p*_spin_ = 0.0001; FDR-corrected) and ventral attention (*p*_spin_ = 0.014; FDR-corrected) networks for HCP-YA and ventral attention (*p*_spin_ = 0.0035; FDR-corrected) and somatomotor (*p*_spin_ = 0.0035; FDR-corrected) networks for HCP-D. Intrinsic networks: vis = visual; sm = somatomotor; da = dorsal attention; va = ventral attention; lim = limbic; fp = fronto-parietal; dmn = default mode.

**Figure S4.**
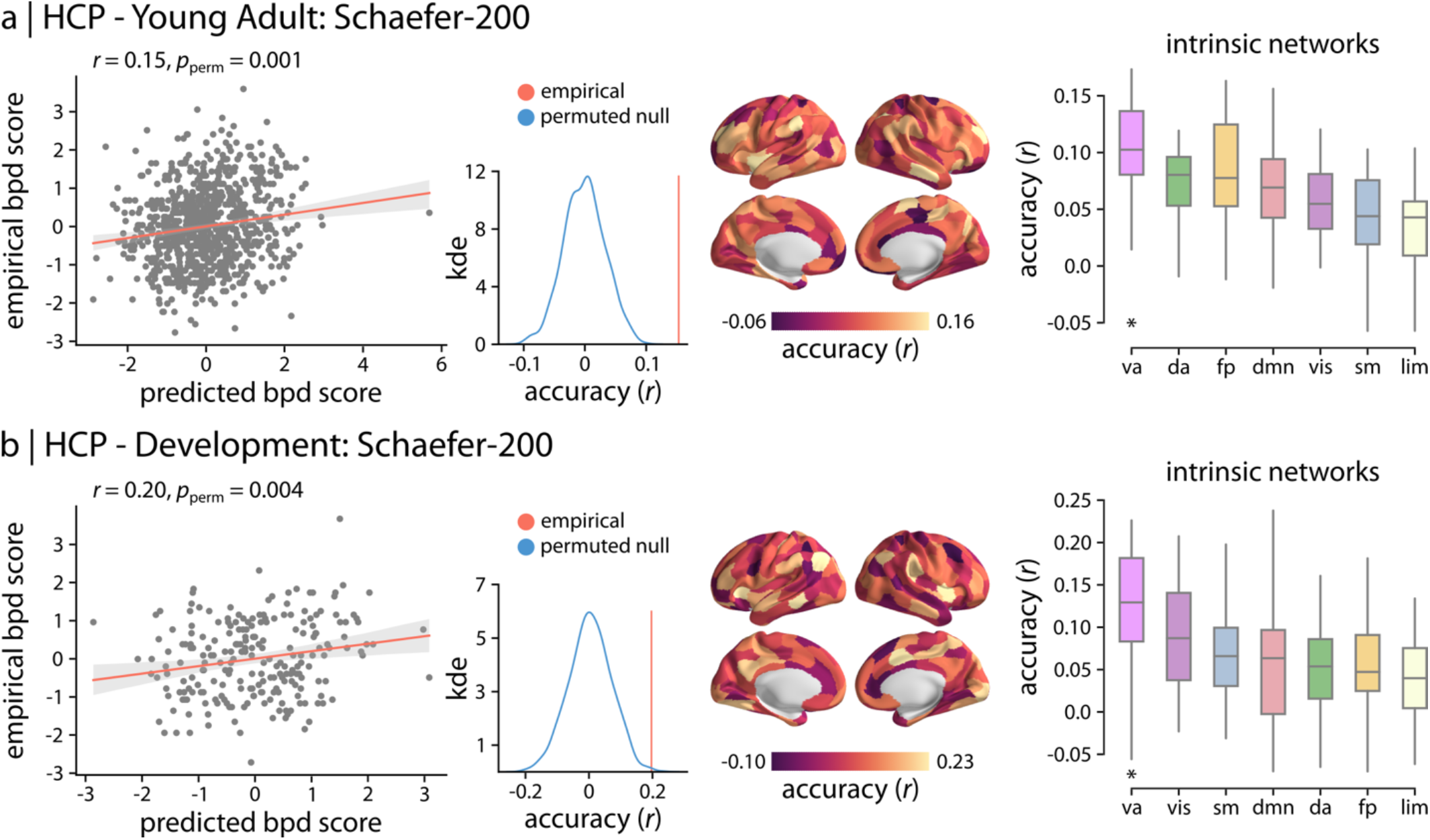
Sensitivity analysis with lower parcellation resolution. To ensure that the findings are independent from the parcellation resolution, a lower resolution atlas with 200 cortical regions (Schaefer-200 atlas) was used to obtain functional connectivity matrices. Functional connectivity data were then used to predict BPD proxy scores. The results are depicted for (a) healthy young adults from the Human Connectome Project (HCP-YA) data and (b) adolescents from the Human Connectome Project - Development (HCP-D). The participant-level prediction accuracies are shown using the scatter plots (HCP-YA: *r* = 0.15, 95% CI = [0.09 0.22]; HCP-D: *r* = 0.20, 95% CI = [0.06 0.32]). Similar to the original analysis, the results were compared with null distributions of accuracies obtained from permutation tests. Region-level accuracies are also depicted across the cortex (Schaefer-400 atlas; 99% confidence intervals). Finally, the average functional system-level prediction accuracy was estimated for the 7 intrinsic functional networks. Asterisk denotes significant system-level prediction accuracy based on 10,000 spatial autocorrelation-preserving null models (*p*_spin_ < 0.05; FDR-corrected). Significant system-level accuracy was observed in ventral attention network for HCP-YA (*p*_spin_ = 0.0001; FDR-corrected) and HCP-D (*p*_spin_ = 0.0001; FDR-corrected). Intrinsic networks: vis = visual; sm = somatomotor; da = dorsal attention; va = ventral attention; lim = limbic; fp = fronto-parietal; dmn = default mode.

**Figure S5.**
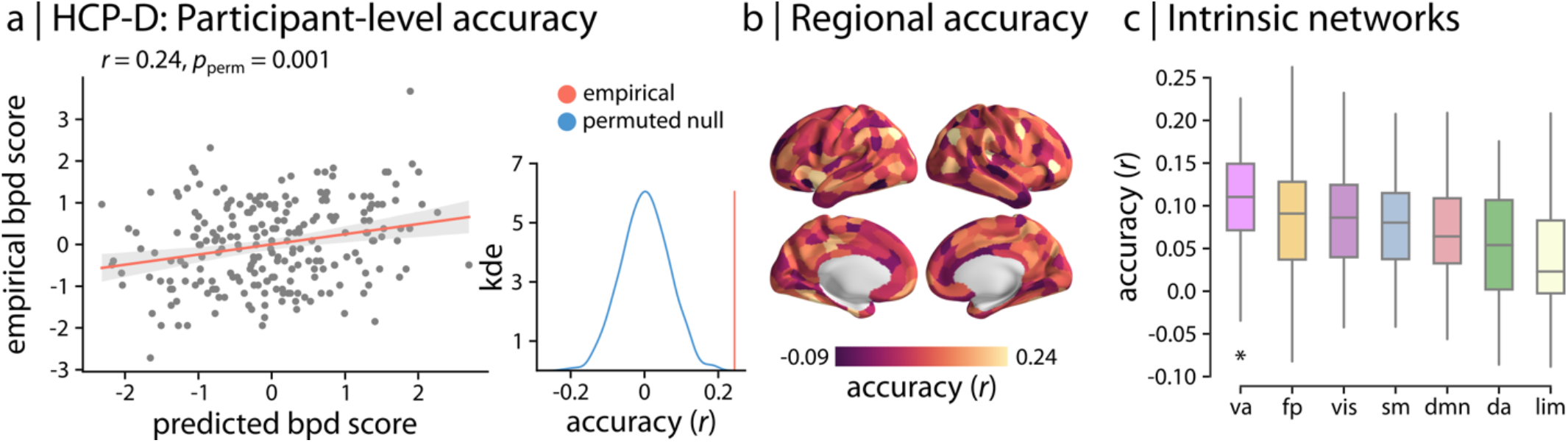
Findings are not influenced by scanning-site effects. We used CovBat-GAM to harmonize functional connectivity across scanning sites in HCP-D and repeated the analyses with harmonized data. Participant-level (a), region-level (b), and system-level (c) accuracies were consistent with the original results. Asterisk denotes significant system-level prediction accuracy (*p*_spin_ < 0.05; FDR-corrected). Significant system-level accuracy was observed in ventral attention network (*p*_spin_ = 0.0001; FDR-corrected). Intrinsic networks: vis = visual; sm = somatomotor; da = dorsal attention; va = ventral attention; lim = limbic; fp = fronto-parietal; dmn = default mode.

**Table S1.** NEO Five Factor Inventory (NEO-FFI) items included in BPD proxy score. The NEO-FFI was used to estimate composite BPD proxy scores (19). The included 24 items and the corresponding questions are listed below. Given that a lower score in items assessing *Agreeableness* and *Conscientiousness* corresponds to more severe BPD symptoms, these items were reverse scored (i.e., multiplied by -1). The composite BPD score was estimated for each participant as the mean score across the included 24 items.

| Neo-FFI Factor | Facet | Item Number and Question |
| --- | --- | --- |
| Neuroticism (N) | Anxiety | #21: "I often feel tense and jittery." |
|  |  | #31: "I rarely feel fearful or anxious." |
|  | Angry Hostility | #36: "I often get angry at the way people treat me." |
|  | Depression | #26: "Sometimes I feel completely worthless." |
|  |  | #41: "Too often when things go wrong, I get discouraged and feel like giving up." |
|  | Vulnerability | #11: "When I'm under a great deal of stress, sometimes I feel like I'm going to pieces." |
|  |  | #51: "I often feel helpless and want someone else to solve my problems." |
| Extraversion (E) | Excitement-Seeking | #22: "I like to be where the action is." |
| Openness (O) | Feelings | #33: "I seldom notice the moods or feelings that different environments produce." |
|  | Actions | #8: "Once I find the right way to do something, I stick to it." |
|  |  | #28: "I often try new and foreign foods." |
| Agreeableness (A) | Trust | #24: "I tend to be cynical and skeptical of others' intentions." |
|  |  | #29: "I believe that most people will take advantage of you if you let them." |
|  | Straightforwardness | #59: "If necessary, I am willing to manipulate people to get what I want." |
|  | Altruism | #14: "Some people think I'm selfish and egotistical." |
|  |  | #39: "Some people think of me as cold and calculating." |
|  | Compliance | #9: "I often get into arguments with my family and co-workers." |
|  |  | #54: "If I don't like people, I let them know it." |
| Conscientiousness (C) | Order | #5: "I keep my belongings clean and neat." |
|  |  | #55: "I never seem to be able to get organized." |
|  | Dutifulness | #40: "When I make a commitment, I can always be counted on to follow through." |
|  |  | #45: "Sometimes I'm not as dependable or reliable as I should be." |
|  | Self-Discipline | #30: "I waste a lot of time before settling down to work." |
|  |  | #50: "I am a productive person who always gets the job done." |

